# NPS neurons receive extensive input from auditory brainstem nuclei

**DOI:** 10.1101/2025.10.08.681223

**Authors:** Richie Zhang, Joel C. Geerling

**Author notes:** Correspondence to: Joel C. Geerling, MD, PhD, PBDB 1320, 169 Newton Rd., Iowa City, IA 52246. Grant sponsors: NIH K08 NS099425 (JCG), NIH R01 NS130038 (JCG), NIH T35 HL166206 (RZ).

## Abstract

Neurons that produce NPS send output to brain regions implicated in circadian function and threat responses, but less is known about the afferent control of NPS neurons. In this study, we used a conventional retrograde tracer, cholera toxin beta subunit (CTb), to identify afferents to the rostral-lateral parabrachial region that contains the main concentration of NPS neurons. We then used Cre-dependent rabies retrograde tracing in *Nps*-2A-Cre mice to identify inputs specifically to NPS neurons. *Nps*-expressing neurons receive heavy input from auditory brainstem structures, including the inferior colliculus, nucleus of the lateral lemniscus, superior olivary complex, and cochlear nucleus. These findings suggest an unexpected role for auditory information in controlling the activity of NPS neurons.

## Introduction

The parabrachial nucleus integrates interoceptive sensory inputs from the hindbrain with descending modulatory inputs from the forebrain to shape behavioral and physiological responses that maintain homeostasis (Carter et al., 2015; Gasparini et al., 2019; Geerling et al., 2016; Grady et al., 2025; Kaur et al., 2017; Mu et al., 2017; Nakamura & Morrison, 2008, 2010; Palmiter, 2018; Shin et al., 2011).

Neurons in this region have a variety of different molecular identities and connectivity patterns (Karthik et al., 2022; Pauli et al., 2022). One subset of neurons at the edge of this region expresses *Nps*, which encodes neuropeptide S (NPS; Adori et al., 2015; Clark et al., 2011; Huang et al., 2022; Xu et al., 2004). Pharmacological studies have shown effects of NPS on locomotion, anxiety, arousal, and food intake (Leonard et al., 2008; Peng et al., 2010; Smith et al., 2006; Xu et al., 2004). Its receptor (*NPSR1*) has been implicated in anxiety, asthma, and endometriosis (Domschke et al., 2011; Laitinen et al., 2004; Tapmeier et al., 2021), and a rare gain-of-function *NPSR1* variant reduces sleep (Xing et al., 2019).

Previously, we described the output projections of *Nps*-expressing neurons in the PB region. These neurons target a variety of brain regions, prominently including the paraventricular nucleus of the thalamus, bed nucleus of the stria terminalis, subparaventricular zone, dorsomedial hypothalamus, and tectal longitudinal column (Zhang et al., 2024). Subsequent studies suggested that *Nps*-expressing neurons in the PB region may promote wakefulness, while those in the central pontine gray may promote sleep (Angelakos et al., 2023; Xing et al., 2024).

The primary concentration of *Nps*-expressing neurons is found in the rostral-lateral PB, dorsal to the Kölliker-Fuse nucleus (Huang et al., 2022), and extends rostrally alongside the lateral lemniscus in the semilunar nucleus (Huang et al., 2022). This population is centered in a subregion described in rats as the “extreme lateral” subnucleus (Fulwiler & Saper, 1984). Unlike other PB subnuclei, the extreme lateral subnucleus does not appear to receive input from viscerosensory neurons in the nucleus of the solitary tract (Herbert et al., 1990), and the sources of input to this rostral-lateral subregion remain unclear.

In this study, we characterized the brain-wide afferents to the rostral-lateral parabrachial region, using conventional retrograde axonal tracing, and to *Nps*-expressing neurons in this region, using rabies retrograde tracing. Our findings revealed an input pattern that suggest an unexpected role for auditory information in controlling the activity of NPS neurons.

## Materials and Methods

### Mice

All mice were group-housed in a temperature- and humidity-controlled room on a 12/12 hr light/dark cycle and with ad *libitum* access to water and standard rodent chow. Overall, we used n=10 female and male mice ranging in age from 9 to 12 weeks (17-29 g body weight). We used *Nps*-2A-Cre and *Nps*-2A-Cre;R26-LSL-L10GFP Cre-reporter mice, with detailed information provided in Table 1. All Cre-driver and reporter mice were hemizygous and maintained on a C57BL6/J background. All experiments were conducted in accordance with the guidelines of the Institutional Animal Care and Use Committee at the University of Iowa.

**Table 1.** Cre-driver and -reporter mice.

| Strain | References | Source Information | Key Gene |
| --- | --- | --- | --- |
| Nps-2A-Cre | Huang, D., Zhang, R., Gasparini, S., McDonough, M. C., Paradee, W. J., & Geerling, J. C. (2022). Neuropeptide S (NPS) neurons: Parabrachial identity and novel distributions. *J Comp Neurol*, 530(18), 3157–3178.<br>https://doi.org/10.1002/cne.25400 | Available from The Jackson Laboratory (Jax #038113) | 2A-Cre inserted immediately after terminal exon in endogenous neuropeptide S gene |
| R26-LSL-L10GFP reporter | Krashes, Michael J., et al. “An excitatory paraventricular nucleus to AgRP neuron circuit that drives hunger.” *Nature* 507.7491 (2014): 238. | Available from originating investigators. | Floxed transcription STOP cassette followed by EGFP/Rpl10 fusion reporter gene under control of the CAG promoter targeted to the Gt(ROSA)26Sor locus |

### Stereotaxic injections

Mice were anesthetized with isoflurane (0.5-2.0%) and placed in a stereotactic frame (Kopf 1900 or 940). We made a midline incision and retracted the skin to expose bregma and the skull over the brain. Through a pulled glass micropipette (20-30 µm inner diameter), we targeted the rostral-lateral parabrachial region in each case, with coordinates: 4.7 mm caudal to bregma, 1.7 mm right of midline, and 3.8 mm deep to bregma. Each injection was made over a 5-minute period using picoliter air puffs through a solenoid valve (Clippard EV 24V DC) pulsed by a Grass stimulator. The pipette was left in place for an additional 3 minutes, then withdrawn slowly before closing the skin with Vetbond (3M). Meloxicam (1 mg/1 kg) was provided for postoperative analgesia.

For non-selective retrograde axonal tracing, we injected cholera toxin beta subunit (CTb; 0.1% in distilled water; List) 30 nL unilaterally in n=5 *Nps*-2A-Cre;R26-LSL-L10GFP Cre-reporter mice. One case (7210) was excluded from analysis because the injection site was tiny and did not produce retrograde labeling.

For Cre-dependent retrograde tracing with modified rabies, we used n=5 *Nps*-2A-Cre mice without a Cre-reporter. First, we made bilateral injections delivering 60 nL of a 1:1 mixture of two “helper viruses” supplying the avian receptor required for entry of EnvA-pseudotyped rabies (AAV8-EF1a-TVA-mCherry; **Table 2**) and the glycoprotein required for G-deleted rabies retrograde entry into non-TVA-expressing afferents (AAV8-CAG-FLEX-RabiesG; **Table 2**). After 14 weeks, we injected 200 nL of EnvA-G-deleted rabies-eGFP, then perfused each mouse 8 days later.

**Table 2.** Antisera used in this study.

| Antigen | Immunogen Description | Source, host species, RRID | Concentration |
| --- | --- | --- | --- |
| Cholera toxin b subunit | Cholera toxin b subunit | List, goat polyclonal, product# 703, lot: 7032A10; RRID: AB_10013220 | 1:10,000 |
| Tyrosine hydroxylase | Denatured TH from rat pheochromocytoma in rabbit host | Millipore, rabbit polyclonal, AB152, 400383, RRID: AB_390204 | 1:10,000 |
| Green fluorescent protein | Full length green fluorescent protein from the jellyfish *Aequorea Victoria* | Thermal Fischer Scientific, Chicken, #A10262, Lot: 1972783, RRID: AB_2534023 | 1:3,000 |
| mCherry | Full length mCherry fluorescent protein | Life Sciences, rat monoclonal, cat. #M11217, lot # R1240561, RRID AB_2536611 | 1:2,000 |

### Perfusion and tissue sections

All mice were anesthetized with a mixture of ketamine-xylazine (i.p. 150-15 mg/kg, dissolved in sterile 0.9% saline), then perfused transcardially with phosphate-buffered saline (PBS), followed by 10% formalin-PBS (SF100-20, Fischer Scientific). After perfusion, brains were removed and fixed overnight in 10% formalin-PBS. We sectioned each brain into 40 µm-thick coronal slices using a freezing microtome and collected tissue sections into separate, 1-in-3 series. Sections were stored in cryoprotectant solution at −20 °C until further processing.

### Immunohistology

For immunofluorescence labeling, we removed the tissue sections from cryoprotectant and rinsed them in PBS before loading them into a primary antibody solution (**Table 3**). Antisera were added to a PBS solution of 0.25% Triton X-100 (BP151-500, Fisher), 2% normal donkey serum (NDS, 017-000-121, Jackson ImmunoResearch), and 0.05% sodium azide (14314, Alfa Aesar) as a preservative (PBT-NDS-azide). We incubated these sections overnight at room temperature on a tissue shaker. The following morning, the sections were washed 3x in PBS and incubated for 2 hours at room temperature in PBT-NDS-azide solution containing species-specific donkey secondary antibodies. These secondary antibodies were conjugated to Cy3, Cy5, or Alexa Fluor 488 (Jackson ImmunoResearch #s 705-165-147, 703-545-155, 711-175-152; each diluted 1:1,000 or 1:500). These sections were then washed 3x in PBS and mounted on glass slides (#2575-plus; Brain Research Laboratories), and coverslipped using Vectashield with DAPI (Vector Labs). Slides were stored in slide folders at 4 °C until imaging.

**Table 3.** Viral vectors.

| Injected Tracer | Origin |
| --- | --- |
| EnvA-G-deleted rabies-eGFP, Titer:<br>2.86*10<sup>8</sup> | Salk Viral Vector Core, RRID:Addgene_32635 |
| AAV8-CAG-FLEX-rabiesG, Titer: 9.2*10<sup>12</sup> | Stanford Vector Core, Lot#: 6751,<br>RRID:Addgene_48333 |
| AAV8-Ef1a-FLEX-TVA-mCherry, Titer:<br>6*10<sup>12</sup> | UNC Vector Core; Lot#: AV5006b,<br>RRID:Addgene_38044 |

### Imaging, analysis, and figures

All slides were scanned using an Olympus VS120 microscope. We began by first acquiring a 2x overview scan then using an 10x objective to scan all tissue sections. We used QuPath (Bankhead et al., 2017) to count cells that contained Rabies-eGFP labeling in all cases. Cell counts are expressed as mean ± standard deviation. We counted every cell that contained in-focus labeling. All counts were reviewed by a senior neuroanatomist (J.C.G.) in conjunction with R.Z. to reach consensus. We used OlyVIA (Olympus) or QuPath to crop full-resolution images and Adobe Photoshop to adjust brightness and contrast. We used Adobe Illustrator to make drawings, plot cells for figures, arrange images, and add lettering for figure layouts. Scale bars were traced in Illustrator atop calibrated lines from cellSens or QuPath to produce clean white or black lines in each figure.

### Nomenclature

For neuroanatomical structures and cell populations, where possible, we use and refer to nomenclature defined in peer-reviewed neuroanatomical literature. In some instances, we use or refer to nomenclature derived from rodent brain atlases (Dong, 2008; Paxinos & Franklin, 2013; Paxinos & Watson, 2007; Swanson, 1992).

## Results

### CTb injection sites

We analyzed retrograde labeling after unilateral CTb injection into the parabrachial region in four *Nps*-2A-Cre;R26-lsl-L10GFP mice. The injection site varied between cases (**Figure 1**). In case 7209, the center of the injection site overlapped the dorsal-medial aspect of the main cluster of NPS neurons in the extreme lateral parabrachial nucleus and extended caudally into the middle of the parabrachial nucleus. In case 7213, the center of the injection site overlapped the dorsal aspect of the main cluster of NPS neurons and extended dorsally into the superior lateral parabrachial nucleus and lateral lemniscus. In case 7216, the center of the injection site overlapped most of the main cluster of NPS neurons and extended dorsally into the lateral lemniscus, ventrally into the Kölliker-Fuse nucleus, and rostrally into the semilunar nucleus. In case 7220, the injection site was centered rostrally in the lateral lemniscus and overlapped the dorsolateral cluster of NPS neurons in the semilunar nucleus. Each injection produced a slightly different pattern of retrograde labeling (**Table 4**), as described below.

**Figure 1.**
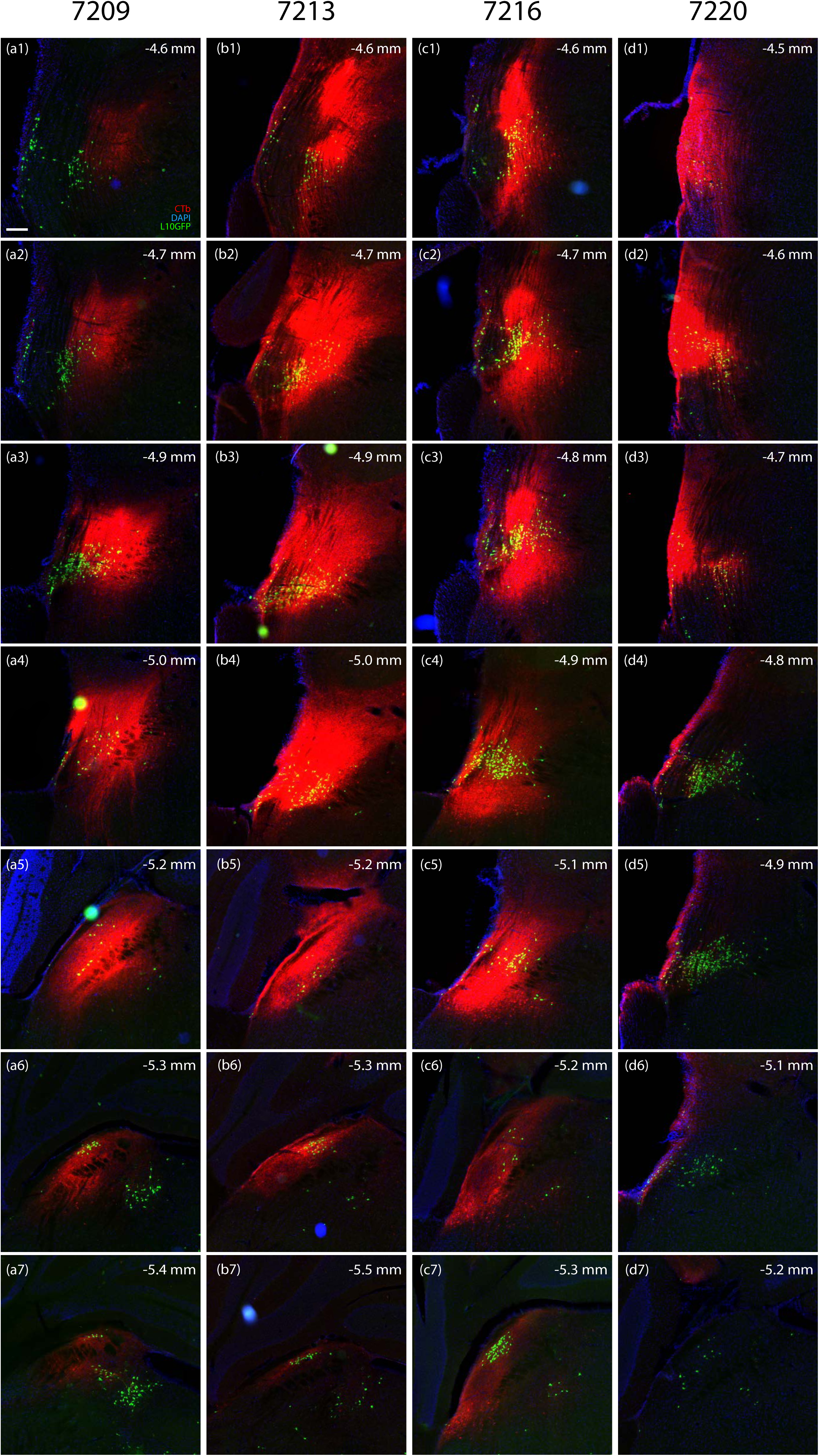
Cholera toxin b subunit (CTb, red) injection sites in *Nps*-2A-Cre;R26-LSL-L10GFP mice (L10GFP in green, DAPI in blue). The CTb injection site is shown in a series of rostral (top row) to caudal (bottom row) levels in each of four cases: (a1-a7) 7209, (b1-b7) 7213, (c1-c7) 7216, and (d1-d7) 7220, with the approximate level caudal to bregma shown in the upper-right corner of each panel. Scalebar in (a1) is 200 µm and applies to all.

**Table 4.**
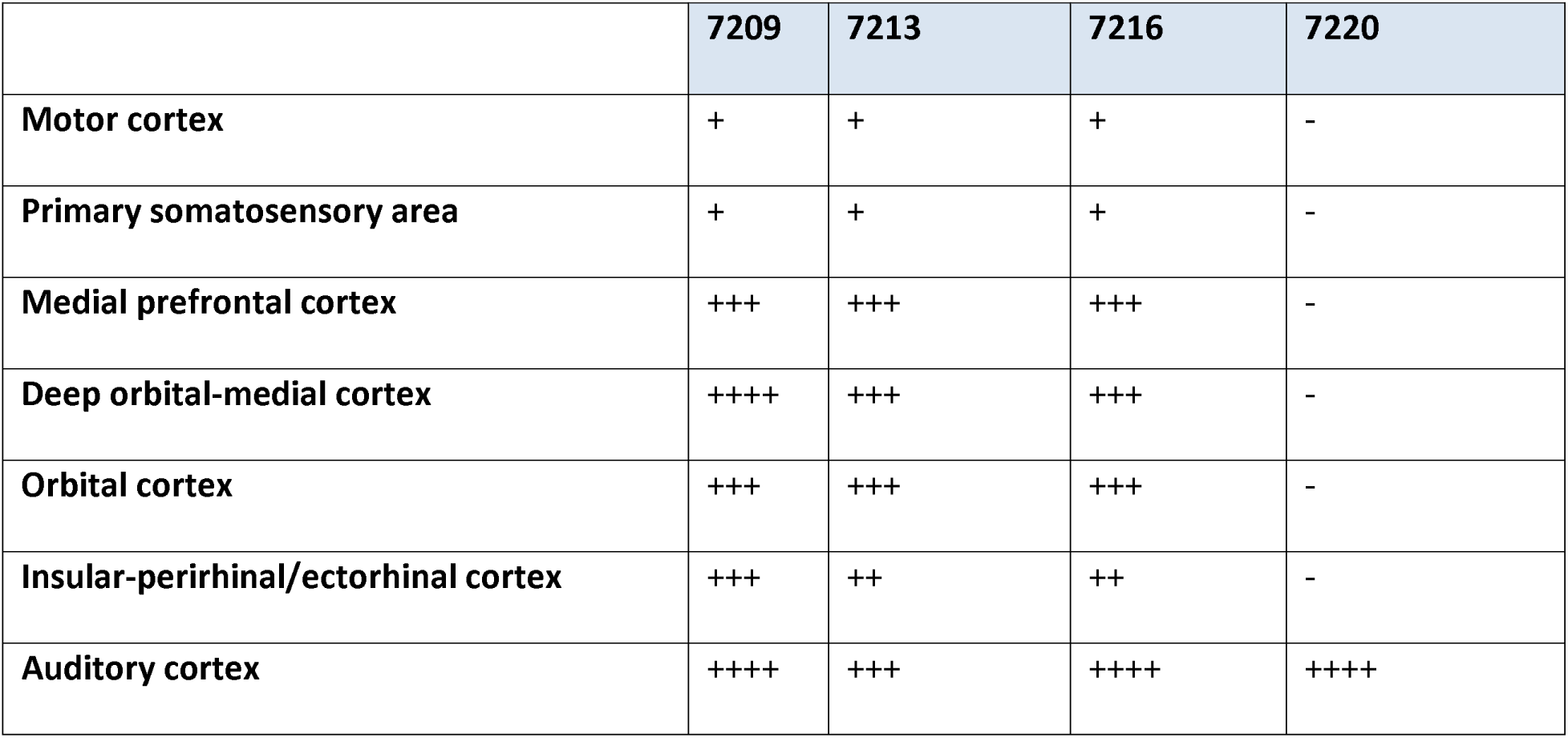
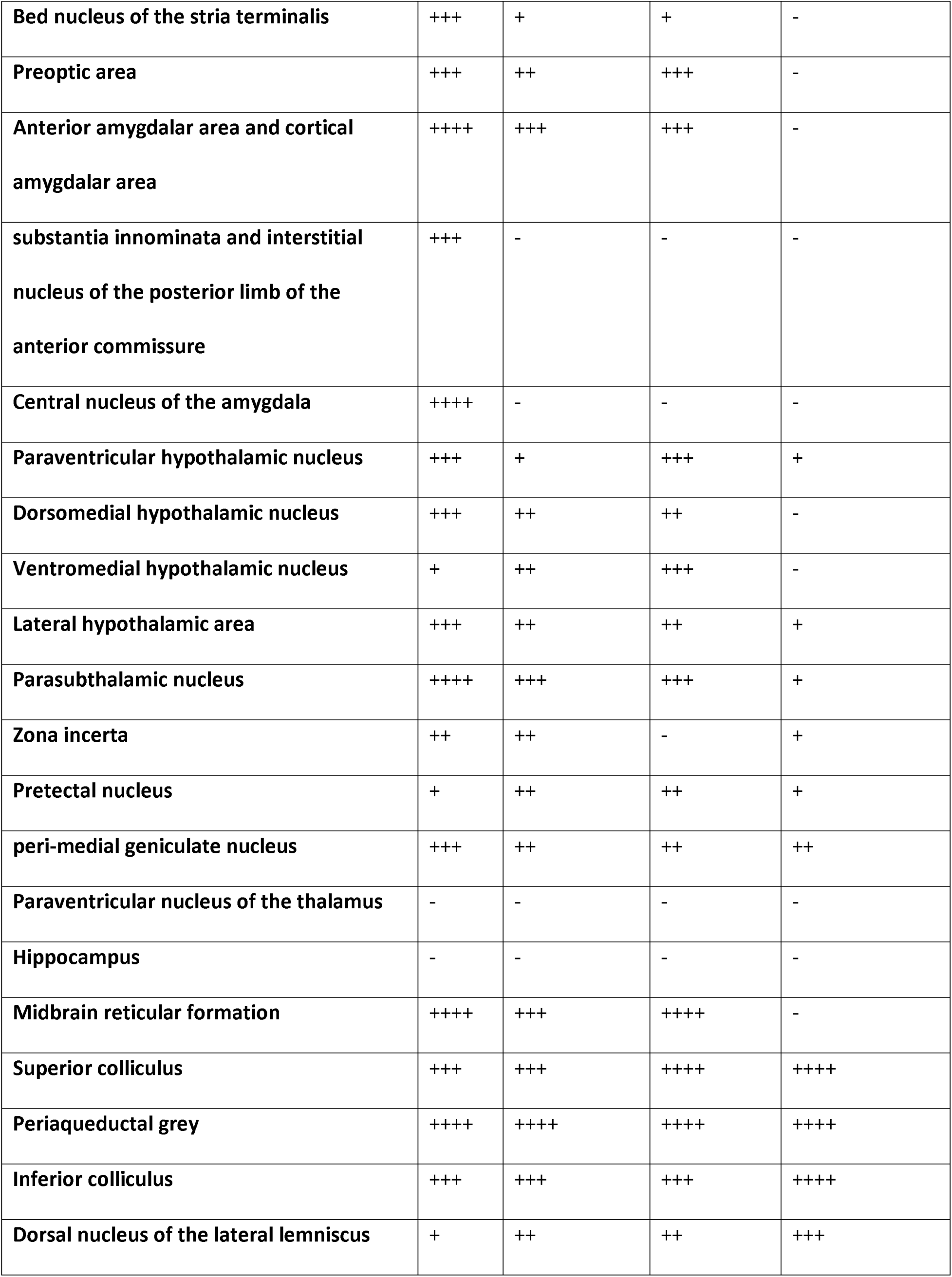
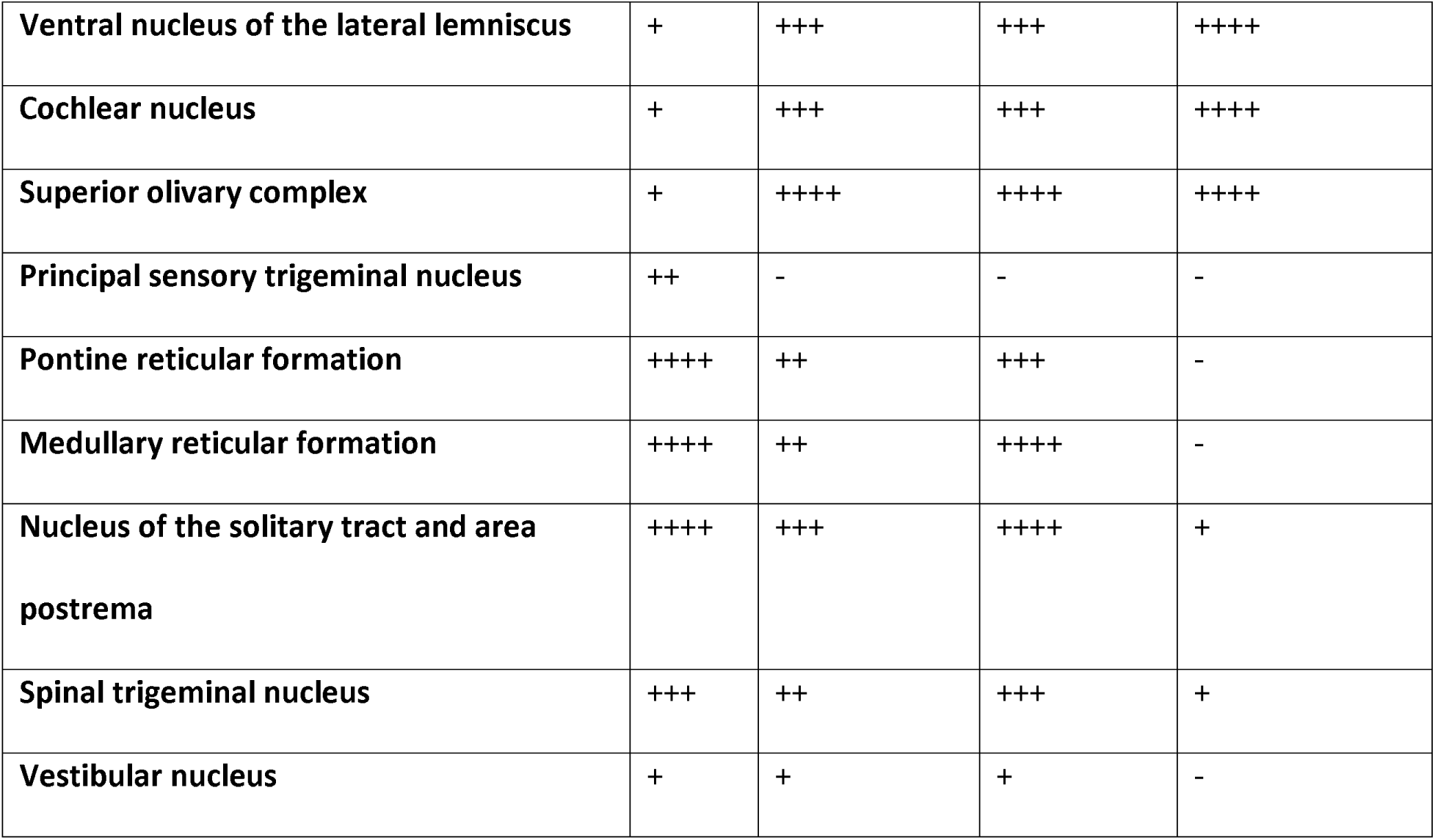
Brain wide ratings of CTb-labeled neurons for all cases. ++++ = 50+ neurons, +++ = 25+ neurons, ++ = 10+ neurons, + = <10 neurons, - = 0 neurons.

### Cerebral cortex

The orbital cortex bilaterally contained a modest number of retrogradely labeled neurons in all cases except 7220. These cases also had many CTb-labeled neurons in the medial prefrontal cortex, within the infralimbic and prelimbic areas. These CTb labeled neurons were present bilaterally but more on the ipsilateral side in every case. Caudally, this retrogradely labeled population extended ventrally to surround the rostro-dorsal pole of the nucleus accumbens (**Figure 2**), occupying a region of the deep orbitomedial prefrontal cortex previously described as “area lambda” (Souza et al., 2022). Each of these cases also had a few CTb-labeled neurons in the ipsilateral primary and secondary motor cortices, and this sparse labeling extended caudally through secondary motor and primary somatosensory cortex. These cases also contained many CTb-labeled neurons in layer 5 of the insular area and in deeper neurons located immediately dorsal to the claustrum and embedded within the external capsule (**Figure 2**) (Grady et al., 2020). Retrograde labeling in this deep cortical region continued caudally into the perirhinal, ectorhinal, and temporal association areas. From there, this pattern of labeling expanded dorsally into layer V of the auditory cortex.

**Figure 2.**
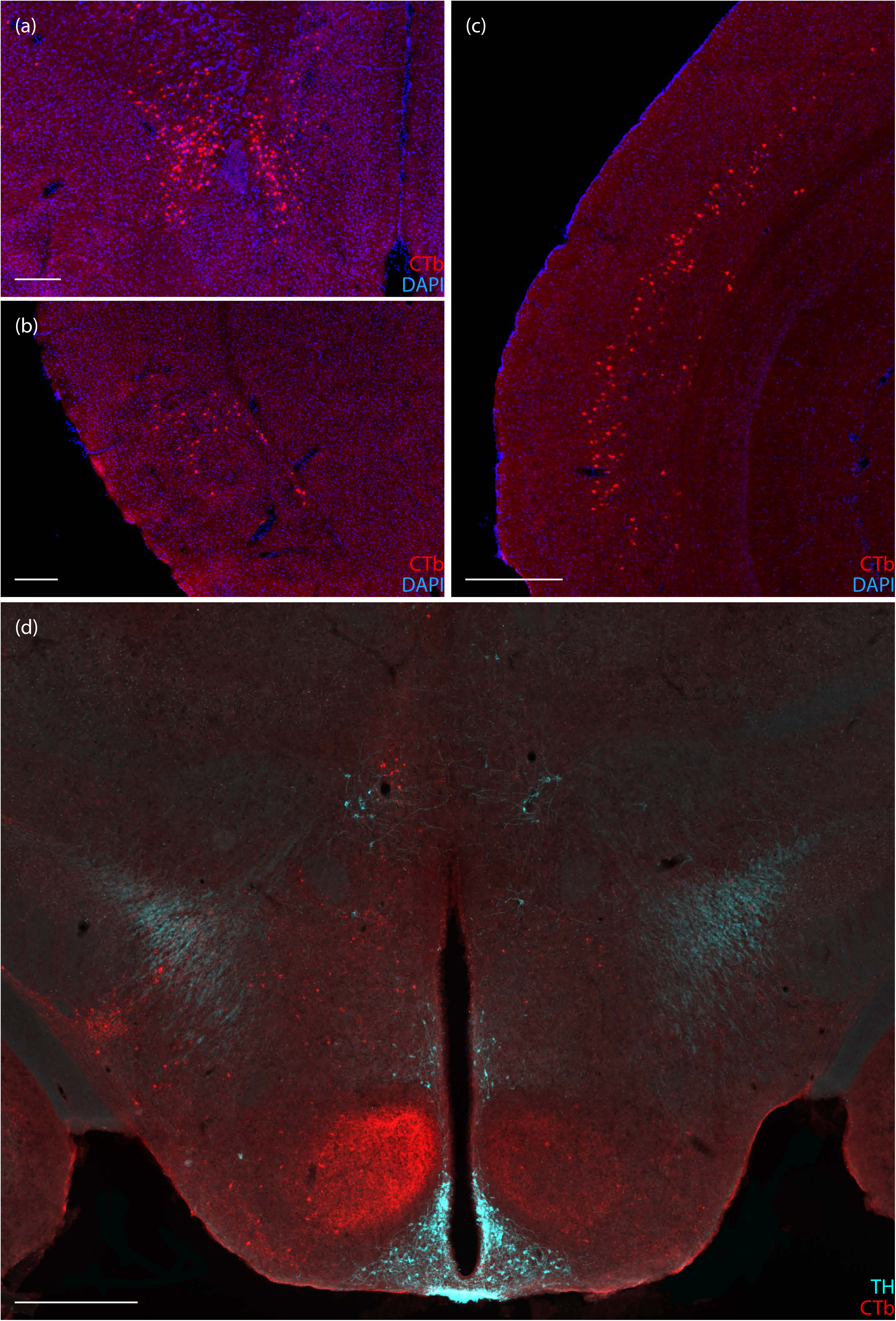
Examples of CTb retrograde labeling in the forebrain. CTb-labeled neurons (red) in the (a) deep orbitomedial prefrontal cortex, immediately rostral-dorsal to the nucleus accumbens (case 7209), (b) insular cortex near the claustrum and embedded in the external capsule (case 7209), (c) auditory cortex (case 7220), and (d) hypothalamus, including anterograde labeling within the VMH (case 7213). CTb-labeled neurons can also been seen above the A11 group in this case, with tyrosine hydroxylase (TH) immunoreactivity shown in ice-blue. Scalebars in (a) and (b) are 250 µm, scalebar in (c) is 400 µm, and scalebar in (d) is 500 µm.

In contrast to the other cases, the pattern of retrograde labeling in the cerebral cortex of case 7220 was shifted caudally and dorsally. While there were no retrogradely labeled neurons in the prefrontal or insular cortex in case 7220, extensive CTb labeling appeared caudally in the auditory cortex (**Figure 2**).

### Bed nucleus of the stria terminalis

While case 7209 had a substantial number of CTb-labeled neurons, primarily in the oval subnucleus, 7216 had only a few labeled neurons in this region, and cases 7213 and 7220 had none.

### Amygdala

Cases 7209, 7213, and 7216 had a moderate number of CTb-labeled neurons in interstitial nucleus of the posterior limb of the internal capsule, plus a few scattered in the cortical amygdalar area. Only case 7209, which extended furthest caudally into the parabrachial nucleus, contained a substantial amount of CTb retrograde labeling in the central nucleus of the amygdala. Cases 7213 and 7216 did not have any labeling in the central nucleus, and 7220 had no labeling in the amygdala.

### Hypothalamus

The medial preoptic area contained a few, scattered neurons in cases 7209, 7213, and 7216 but none in case 7220. While all cases had CTb-labeled neurons in the paraventricular hypothalamic nucleus, case 7209 had the most. We found light labeling scattered across several other hypothalamic nuclei (including lateral, dorsomedial, arcuate, and ventromedial), more in case 7209 than others. Case 7209 also had many CTb-labeled neurons in the parasubthalamic nucleus, which had light to moderate labeling in cases 7213, 7216, and 7220.

Although anterograde labeling was not a focus of our study, the pattern in case 7213 was remarkable for prominent CTb labeling in axons in the dorsomedial part of the ventromedial hypothalamic nucleus (**Figure 2**), reflecting a previously identified heavy projection from the superior lateral parabrachial subnucleus (Bester et al., 1997; Huang et al., 2020; Karthik et al., 2022). Cases 7209 and 7216 had light labeling here, but 7220 lacked anterograde labeling in the ventromedial nucleus.

### Thalamus

The medial geniculate complex was surrounded by many retrogradely labeled neurons in case 7216, but this region contained few neurons in cases 7213 and 7209 and only light axonal labeling in case 7220. In cases 7213, 7216, and 7220, but not 7209, we found a cluster of retrogradely labeled neurons in the caudal paramedian thalamus, near the A11 group of dopaminergic neurons, in the location of the magnocellular subparafasicular thalamic nucleus (**Figure 3**). Case 7213 had CTb-labeled neurons in the posterior limitans nucleus. Also, the pretectal region contained labeling in each case.

**Figure 3.**
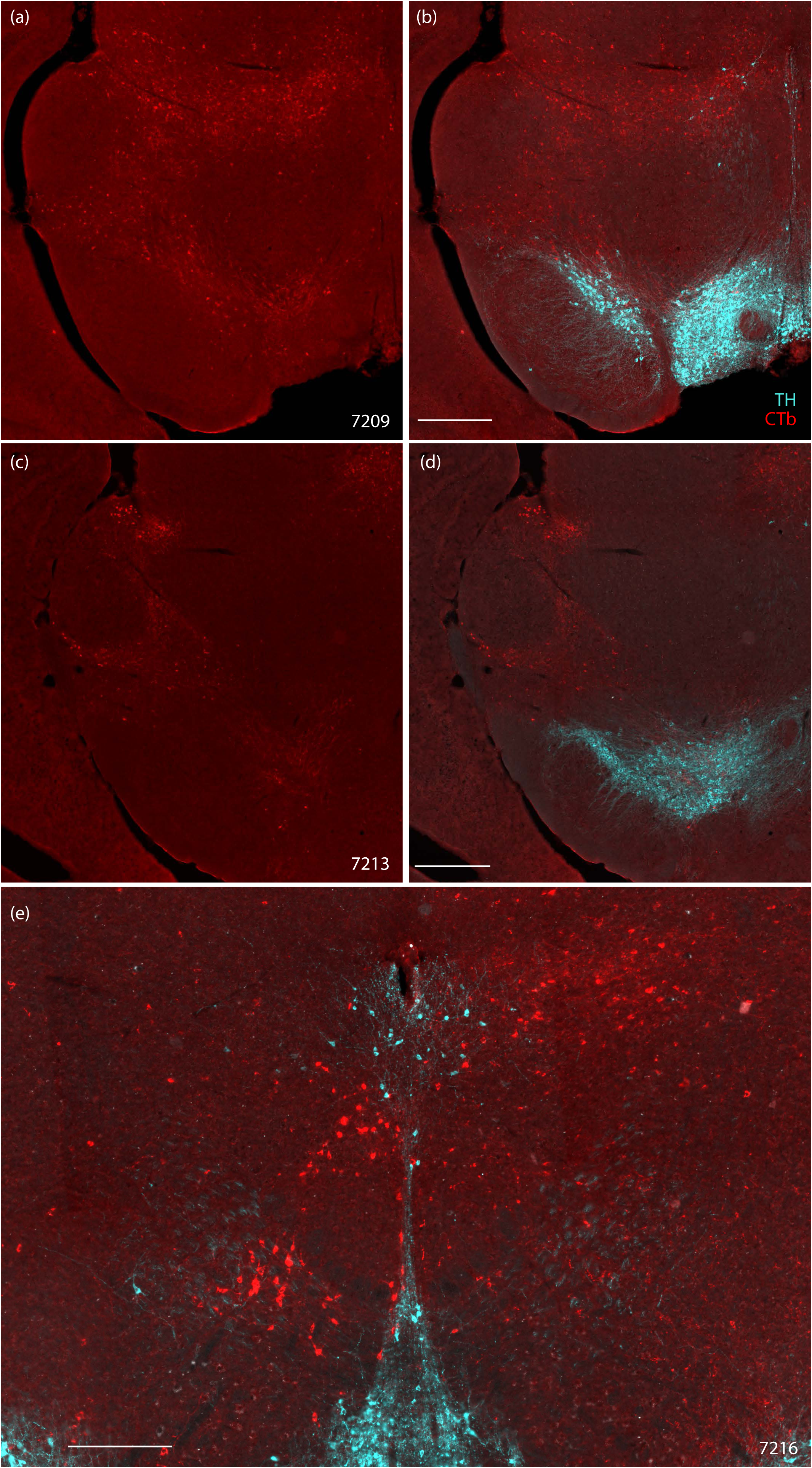
CTb-labeled neurons (red) in the midbrain reticular formation in (a) case 7209, (c) case 7213, shown with TH immunoreactivity (ice-blue) in (b) and (d) respectively. CTb-labeled neurons surrounding the oculomotor nucleus in case 7216 with TH immunolabeling (ice-blue). Scalebar in (b) is 500 µm and applies to (a). Scalebar in (d) is 500 µm and applies to (c). Scalebar in (e) is 250 µm.

### Midbrain

In the superior colliculus, both intermediate and deep layers contained many neurons in case 7216, largely on the ipsilateral side. In comparison, cases 7209 and 7213 had moderate amount of labeling in the superior colliculus. Case 7220 had dense CTb-labeling in both deep and superficial layers of the superior colliculus.

There were many CTb-labeled neurons in the periaqueductal grey in every case but the distribution differed between cases. In case 7209, CTb-labeled neurons were distributed in the ventrolateral column; in case 7213, in the dorsolateral column; in case 7216, in both the dorsal and ventrolateral columns; and in case 7220, the majority were located in the dorsal column, with a few in the ventrolateral column.

In the ventral midbrain, a moderate number of CTb-labeled neurons spanned the retrorubral field, substantia nigra pars compacta, and ventral tegmental area in case 7209, fewer in case 7213, and none in cases 7216 and 7220. CTb labeling was intermingled but mutually exclusive with neurons that were immunoreactive for tyrosine hydroxylase, which is a marker of dopaminergic neurons in this region.

The midbrain reticular formation contained different patterns across cases (**Figure 3**). A moderate number of CTb-labeled neurons surrounded the medial geniculate nucleus in case 7213 and case 7216. None were found in this area in case 7220. Case 7209 had many CTb-labeled neurons spread throughout the midbrain reticular formation surrounding the red nucleus and extending along the dorsal aspect of the substantia nigra and ventral tegmental area. Case 7216 had scattered labeling in the midbrain reticular formation and prominent labeling surrounding the contralateral oculomotor nucleus, including CTb labeled neurons intermingled with TH-immunoreactive axons in the dorsal noradrenergic bundle.

In the caudal midbrain, every case had retrograde labeling in the inferior colliculus, with a wide range of labeling density and intensity across cases. Retrograde labeling in the inferior colliculus was light in 7209 (caudal injection site, primarily in the parabrachial nucleus) but very dense in 7220 (rostral injection site, primarily in the lateral lemniscus), with intermediate labeling densities in 7213 and 7216. In each case, retrogradely labeled neurons in the inferior colliculus were distributed primarily in the dorsal and external cortex, outside the central nucleus.

### Hindbrain

The contralateral cochlear nucleus contained many CTb-labeled neurons in case 7220. We also found a moderate amount of labeling in cases 7213 and 7216 but none in 7209. The superior olivary complex had many CTb-labeled neurons bilaterally in case 7216 and 7213, prominently ipsilateral labeling in 7220, and none in 7209.

The reticular formation contained a moderate number of CTb-labeled neurons in cases 7209 and 7216 but none in cases 7213 and 7220. Cases 7216 and 7209 also had many CTb-labeled neurons in the nucleus of the solitary tract and area postrema, but this region contained only a few neurons in cases 7220 and 7213.

Case 7216 (injection site extended ventrally, into the Kölliker-Fuse nucleus) had a prominent concentration of CTb-labeled neurons in the ventrolateral medulla, all of which were mutually exclusive with the tyrosine-hydroxylase-immunoreactive neurons of the A1 noradrenergic cell group in this region. This region contained fewer, scattered neurons in cases 7209 and 7213. Case 7220 had no neurons in the caudal ventrolateral medulla and instead had a smaller cluster of CTb-labeled neurons between the inferior olive and the rostral ventrolateral medulla.

The spinal trigeminal nucleus contained a moderate number of CTb-retrogradely labeled neurons in cases 7209 and 7216, with fewer in 7213 and only sparse labeling in 7220. A moderate amount of CTb-labeled neurons were found rostral and dorsolateral to the facial motor nucleus, flanking cranial nerve VII, in the pontine reticular formation of case 7209. In contrast, case 7216 contained prominent anterograde labeling in the facial motor nucleus and scattered neurons in the pontine reticular formation. Cases 7213 and 7220 had very few neurons in this region.

### Rabies injection sites

We made bilateral injections targeting the rostral parabrachial region with Cre-conditional rabies retrograde tracing in five *Nps*-2A-Cre mice. Injection sites are shown in **Figures 4** (case 6273) and **5** (cases 6269-6272). Three cases (6269, 6270, and 6272) had unilateral and two cases (6271 and 6273) had bilateral “starter cells,” which co-expressed TVA-mCherry and rabies eGFP. Unilateral case 6269 had starter cells and neurons expressing TVA-mCherry alone on the left side only. Unilateral cases 6270 and 6272 had TVA-mCherry-expressing neurons on both sides and rabies starter cells on the right side only. The pattern of rabies eGFP retrograde labeling was highly similar across cases. Labeling was similar between left-side (6269) and right-side (6270 and 6272) cases, and there were no left-right differences in the labeling pattern in either of the bilateral cases (6271 and 6273). We selected case 6273, which had the most rabies starter cells bilaterally, as the representative case to display this common brain-wide labeling pattern (**Figure 6**).

**Figure 4.**
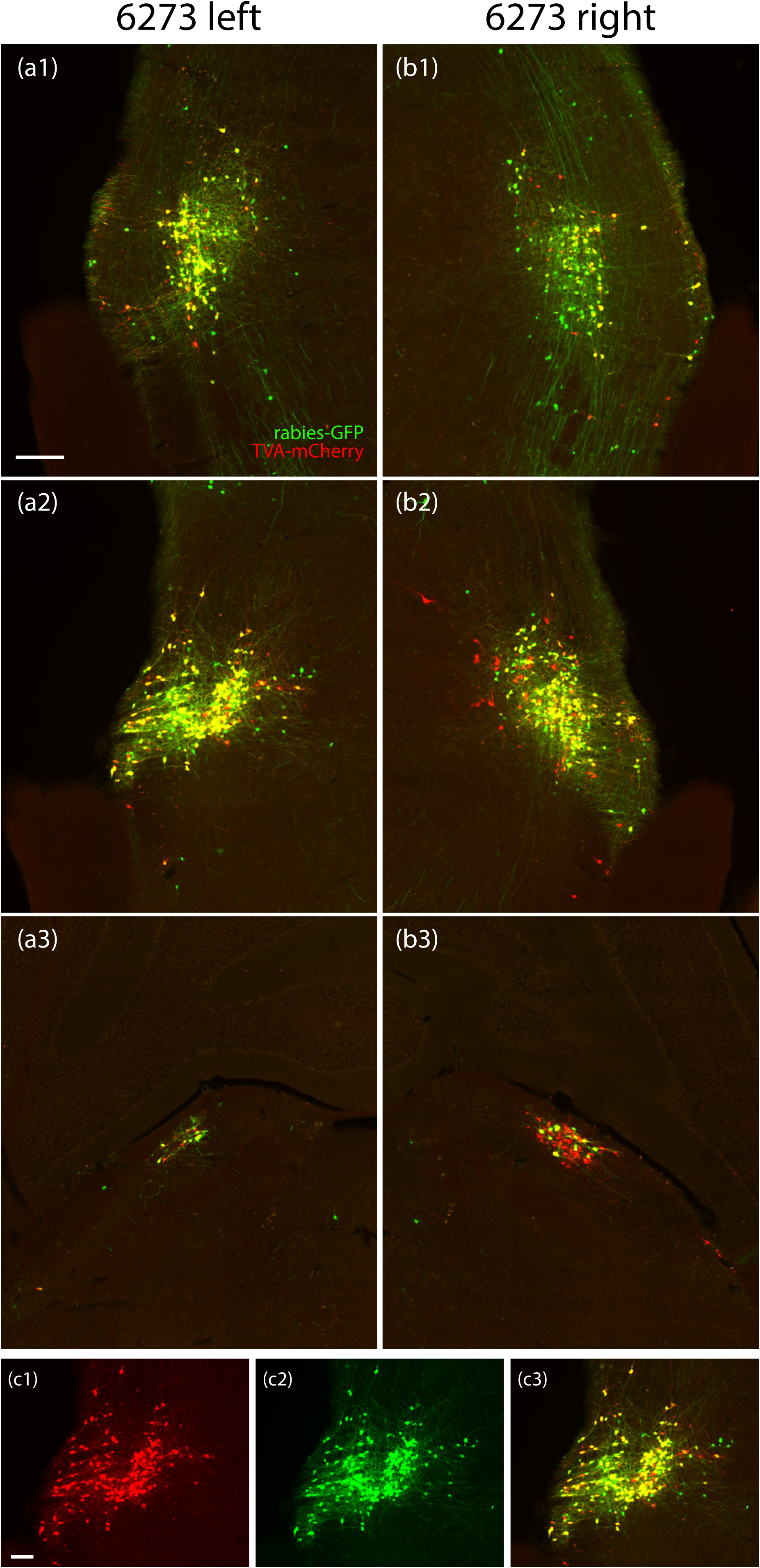
Rabies injection sites for case 6273, which was used for the brain-wide plot. Neurons co-labeled with rabies-eGFP (green) and TVA-mCherry (red) are shown on both sides of the brain at three rostrocaudal levels through the injection site: a rostral section including the semilunar nucleus on the left (a1) and right (b1); a middle section including the extreme lateral parabrachial nucleus on the left (a2) and right (b2); and a caudal section including the dorsolateral parabrachial nucleus on the left (a3) and right (b3). Color separation showing TVA-mCherry (c1), rabies-eGFP (c2), and both (c3) in the left-side extreme lateral parabrachial nucleus from panel (a2). Scalebar is 200 µm in (a1) and applies to (a2-a3, b1-b3). Scalebar is 100 µm in (c1) and applies to (c2) and (c3).

**Figure 5.**
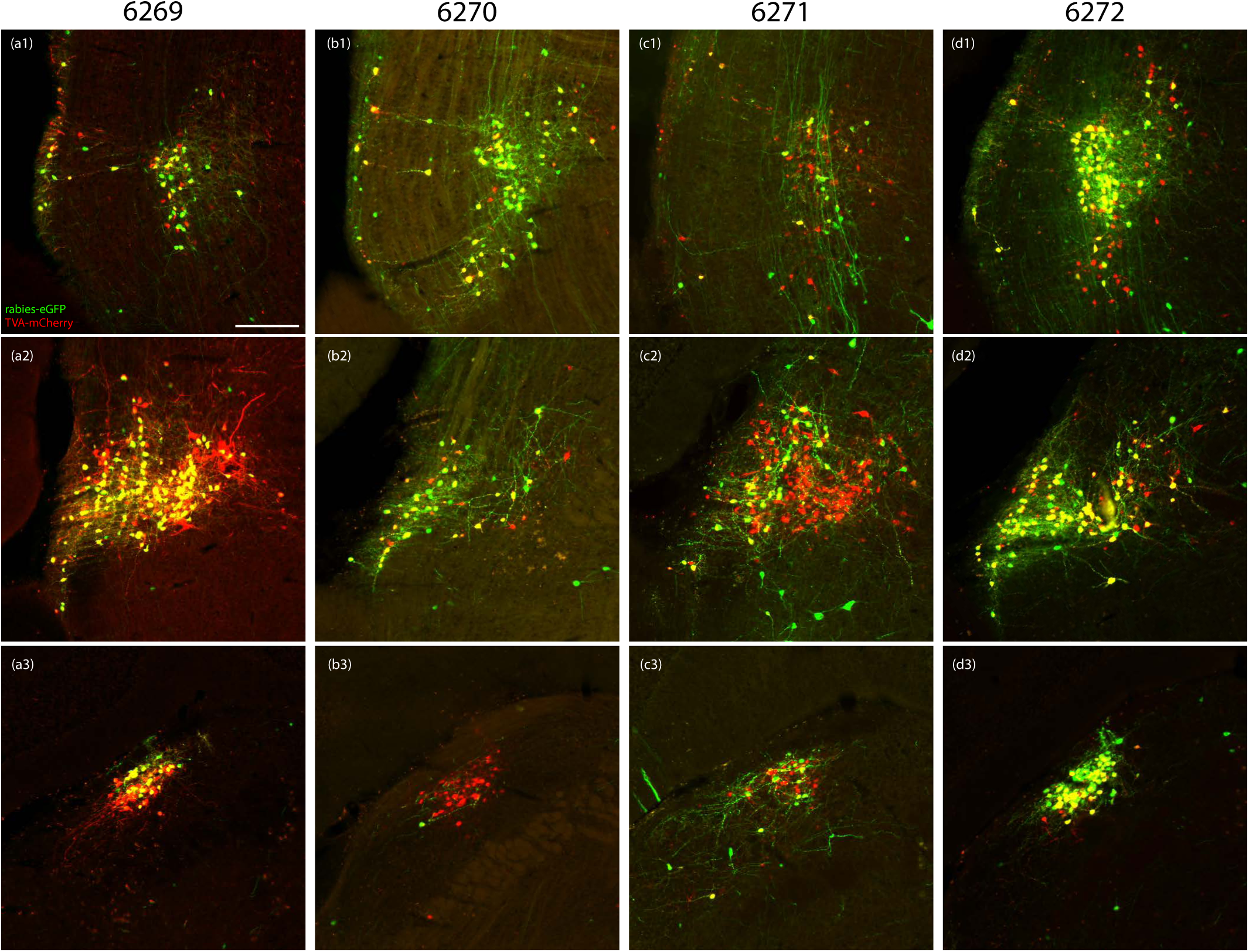
Rabies injection sites for all remaining cases. Rabies-eGFP (green), TVA-mCherry (red), and co-labeled “starter cells” (yellow) are shown for case 6269 (a1-a3, left), 6270 (b1-b3, right), 6271 (c1-c3, right), and 6272 (d1-d3, right). For side-by-side comparison, every injection site is shown on the left side. The contralateral injection site in bilateral case 6271 is not shown; the other three cases had starter cells only on the side shown in this figure. Scalebar in (a1) is 200 µm and applies to all.

**Figure 6.**
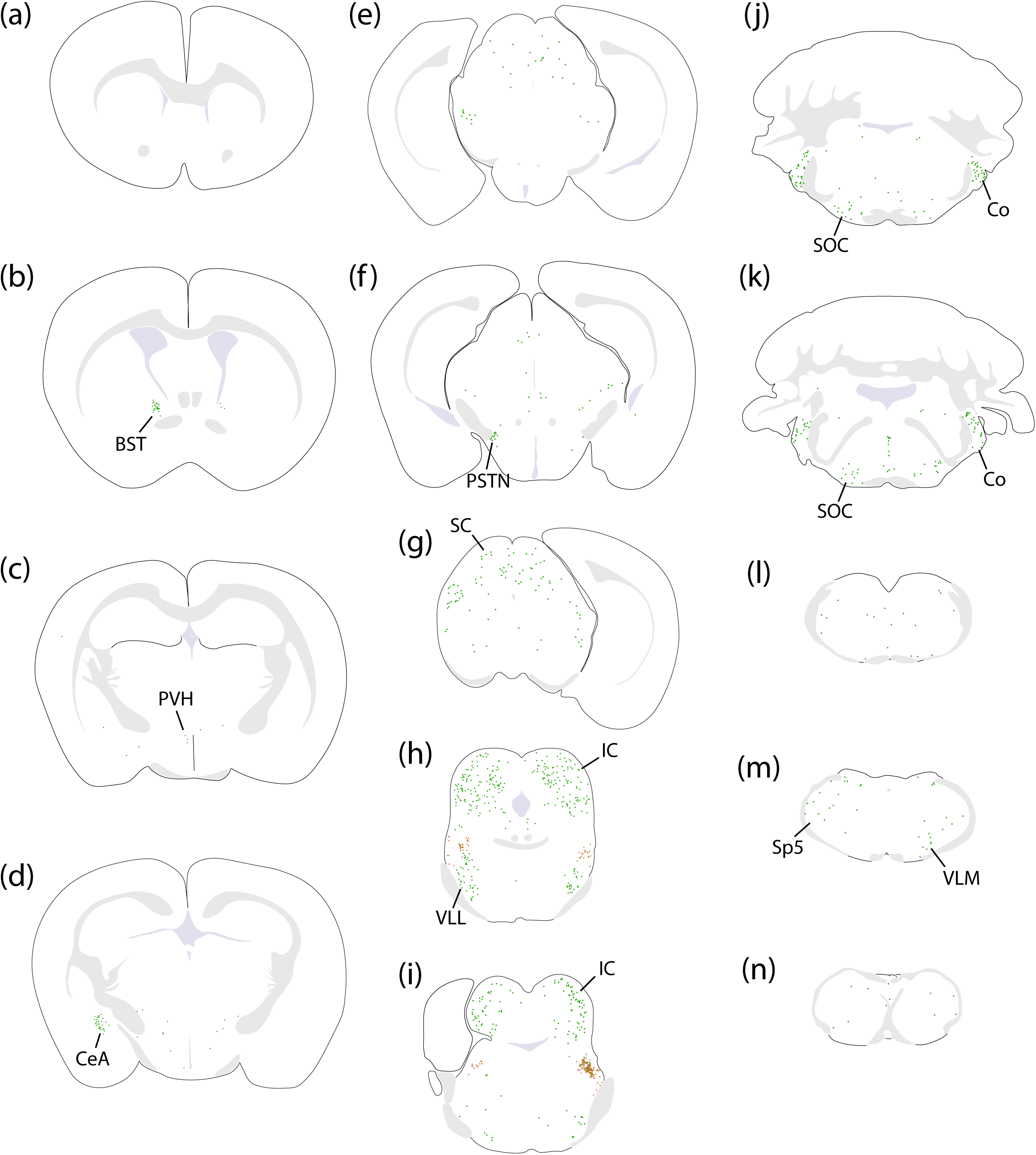
Rabies retrograde labeling across the brain (a-n) in a representative case with bilateral injection sites (6273). Each green dot represents a rabies-eGFP-expressing neuron. In panels (h) and (i), “starter cells” expressing both rabies-eGFP and TVA-mCherry are shown as a red ring around a green dot, and neurons expressing just TVA-mCherry are shown as light-red dots.

### Cerebral cortex

In contrast to the extensive cortical labeling in CTb cases, only a few rabies retrogradely labeled neurons were seen in the cerebral cortex and orbital cortex, only in bilateral cases 6271 and 6273. These neurons had a morphology characteristic of pyramidal neurons (**Figure 7**).

**Figure 7.**
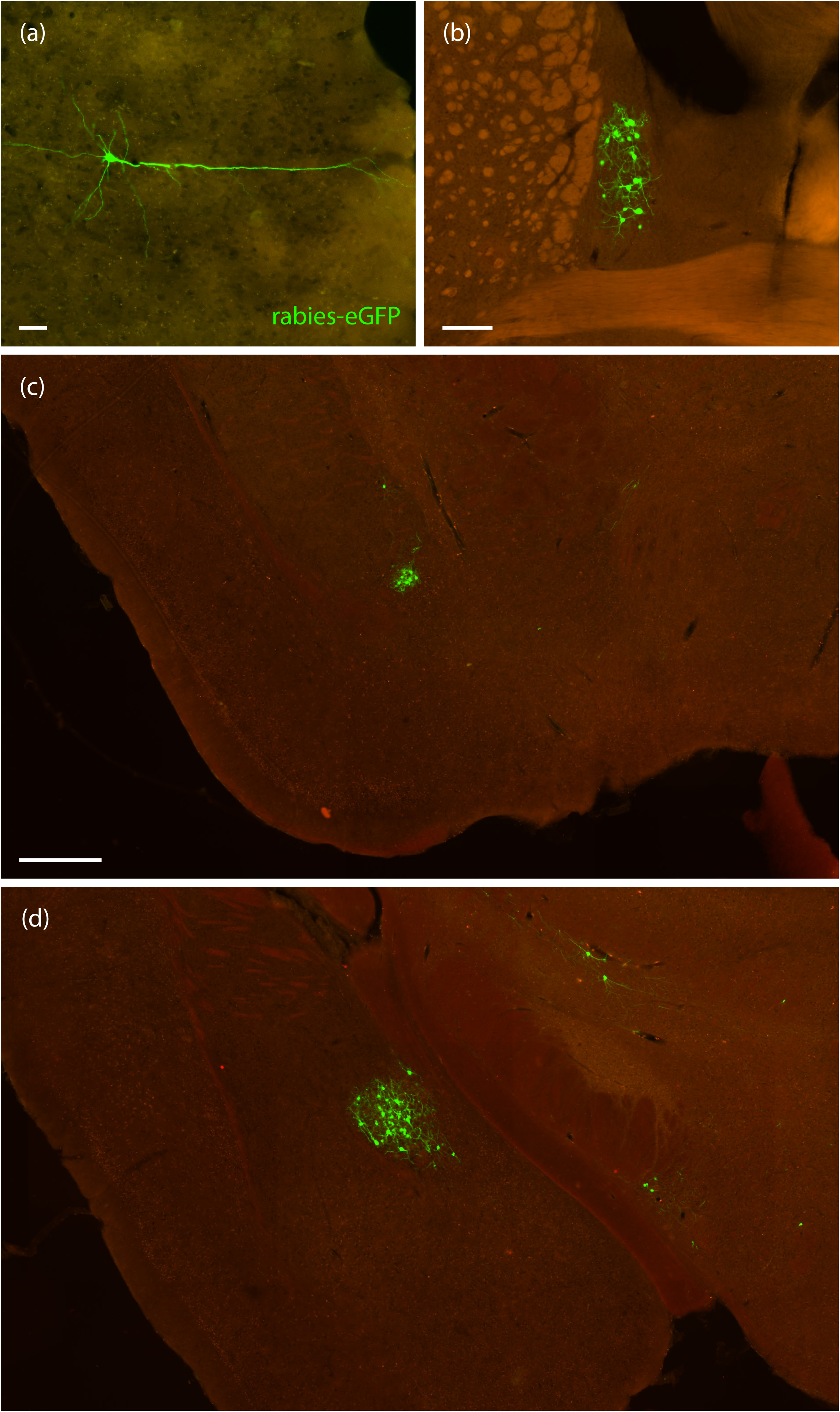
(a) Rabies-eGFP-labeled (green) pyramidal neuron in the perirhinal cortex of case 6271. (b) Rabies-eGFP-labeled neurons in the dorsal bed nucleus of the stria terminalis of case 6270. (c) Cluster of rabies-eGFP-labeled neurons in the interstitial nucleus of the posterior limb of the internal capsule and (d) central nucleus of the amygdala in case 6273. Scalebar in (a) is 50 µm. Scalebar in (b) is 200 µm. Scalebar in (c) is 400 µm and applies to (d).

### Bed nucleus of the stria terminalis

The bed nucleus of the stria terminalis contained rabies retrogradely labeled neurons in every case. Virtually all of these neurons were in the oval subnucleus, but we found an individual neuron ventral to the anterior commissure in three cases (**Figure 7**).

### Amygdala

A dense cluster of rabies retrogradely labeled neurons were seen in the interstitial nucleus of the posterior limb of the internal capsule in every case (**Figure 7**). Caudally, a larger group of rabies retrogradely labeled neurons was seen in the central nucleus of the amygdala, primarily in its lateral subdivision, with occasional neurons in the capsular and medial subdivisions (**Figure 7**).

### Hypothalamus

No cases contained prominent labeling in the hypothalamus. Rostrally, a few neurons were found in the preoptic area, and each case contained a small number of neurons in the paraventricular hypothalamic nucleus. We also found a handful of neurons scattered across the lateral hypothalamic area in each case. Between the thalamus and hypothalamus, we found a consistent, moderate number of rabies-GFP-labeled neurons in the parasubthalamic nucleus and zona incerta in every case.

### Thalamus

The thalamus contained very few rabies retrogradely labeled neurons. None of the principal or intralaminar thalamic nuclei contained labeling in any case. We only found an occasional neuron in the pre-tectal region, just rostral to the superior colliculus.

### Midbrain

Auditory-related regions of the midbrain contained the greatest numbers of neurons in every case, both in absolute count (**Figure 8**) and in proportion to the total number of neurons labeled in each brain (**Figure 9**). The inferior colliculi contained the greatest number of rabies-GFP labeled neurons in every case. In the rostral inferior colliculus, at the level where the superior colliculi disappear, rabies-GFP labeling appeared to concentrate in three layers, forming a tulip shape (**Figure 10**). Caudally, labeled neurons avoided the central nucleus and concentrated mostly in the lateral cortex of the inferior colliculus (**Figure 10**). Also, the ventral nucleus of the lateral lemniscus contained many rabies-GFP labeled neurons in every case. In comparison, the dorsal nucleus of the lateral lemniscus contained fewer neurons.

**Figure 8.**
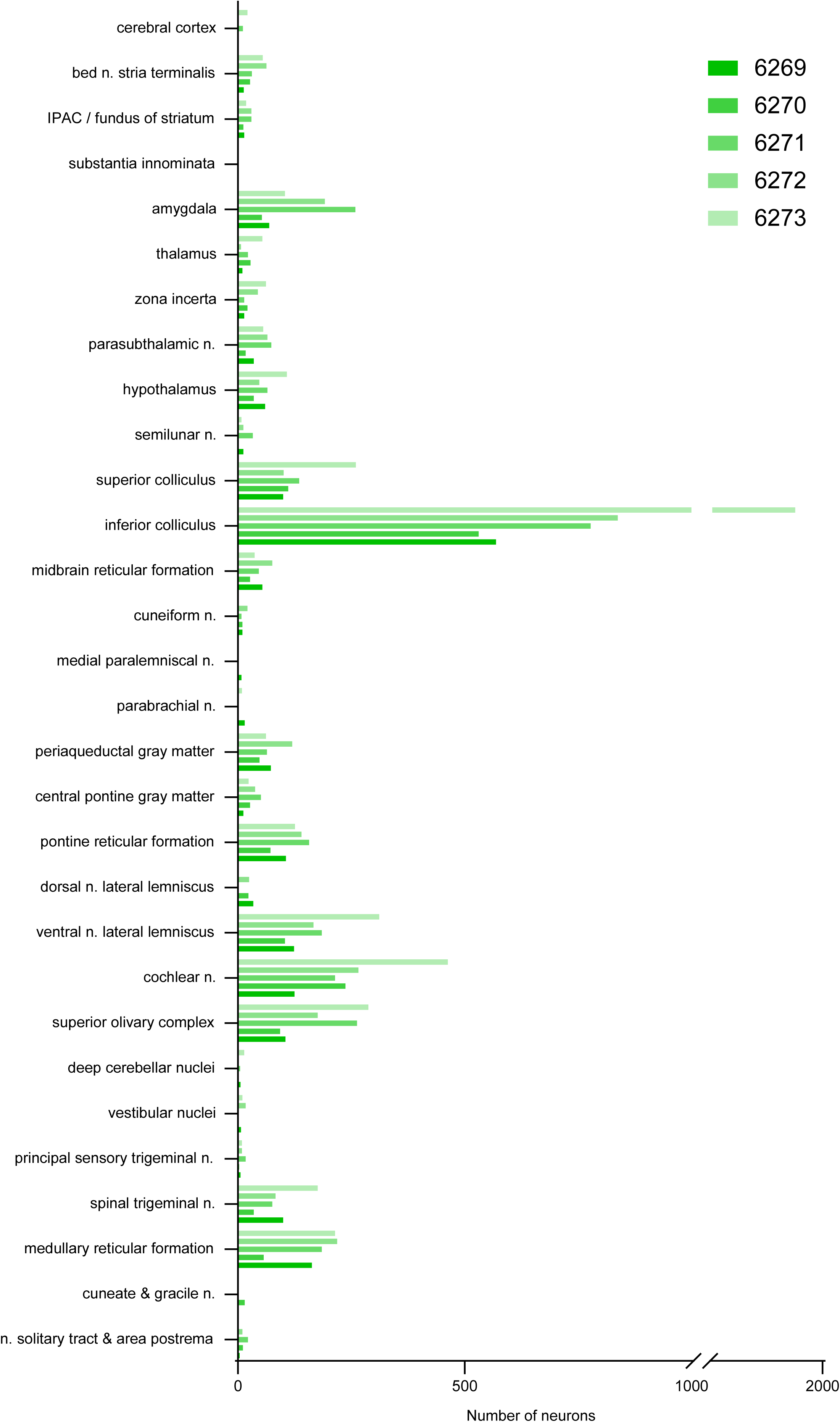
Total counts of rabies-eGFP-labeled neurons, separated by brain region and case number.

**Figure 9.**
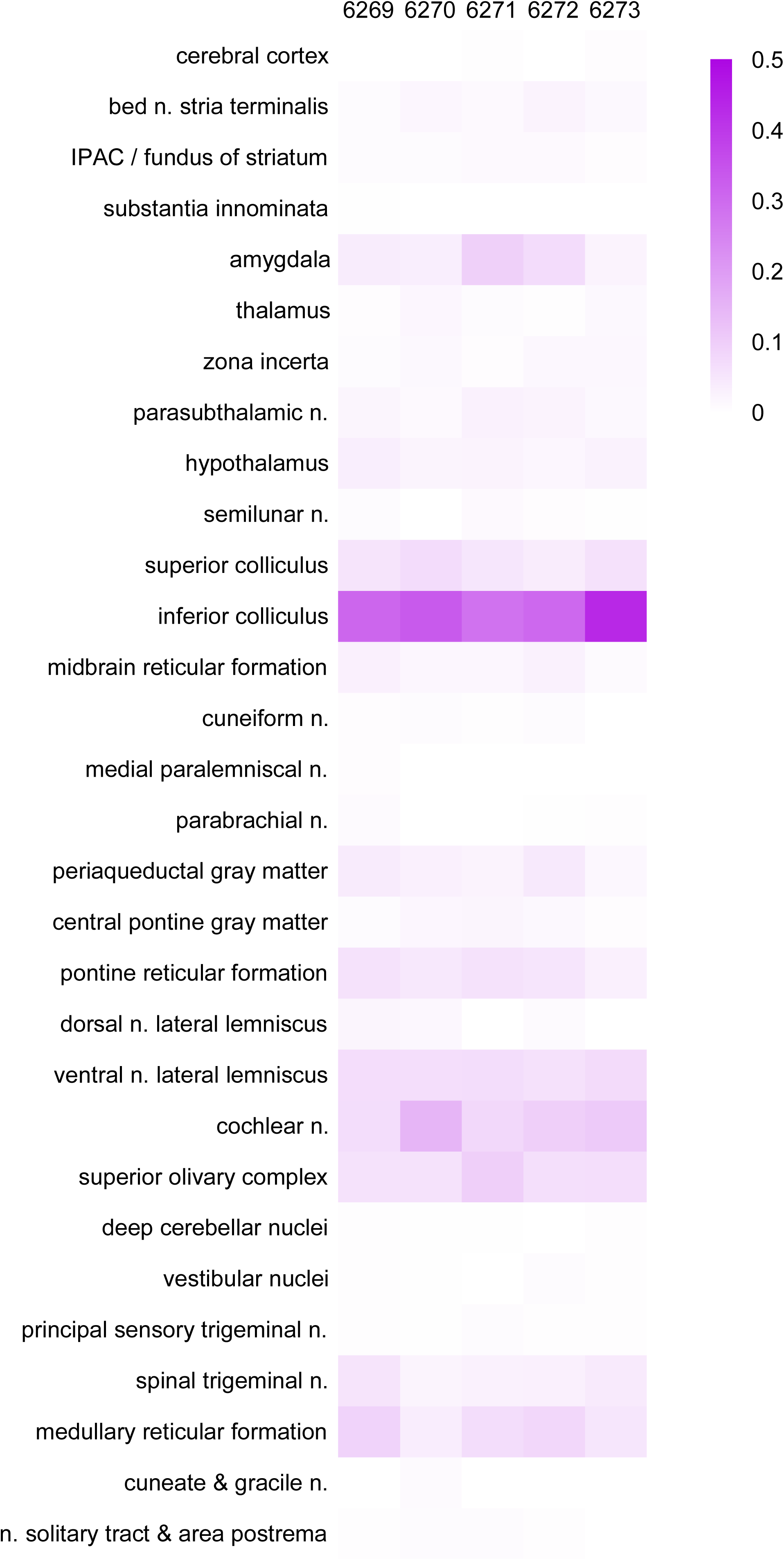
Heatmap showing rabies-eGFP-labeled neurons across brain regions as a proportion of total neurons counted in each case.

**Figure 10.**
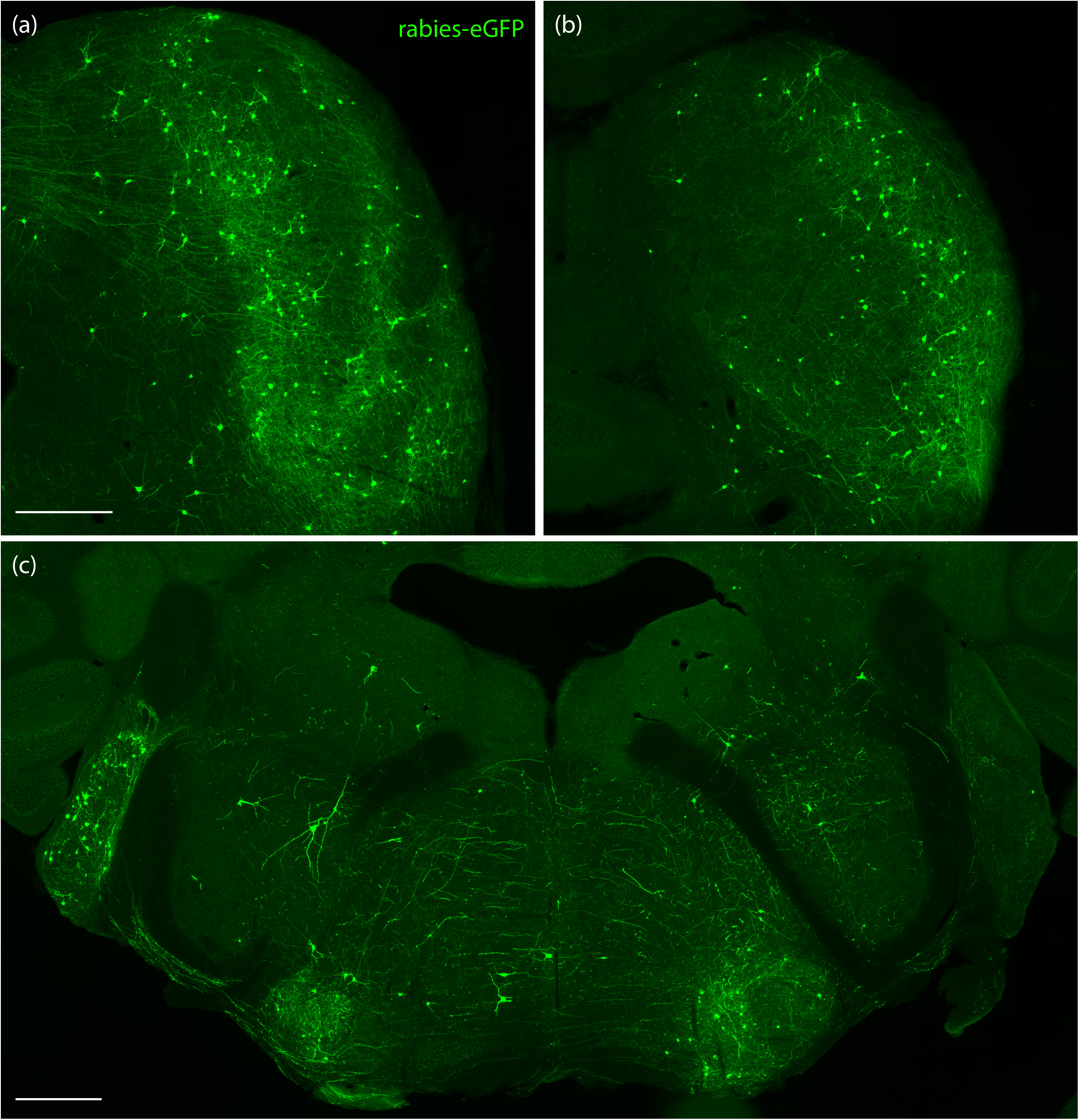
Representative rabies-eGFP (green) labeling pattern in the rostral (a) and caudal (b) inferior colliculus of case 6273. (c) Rabies-eGFP labeled neurons in the cochlear nucleus, superior olivary nucleus, and reticular formation of case 6272. Scalebar in (a) is 400 µm and applies to (b). Scalebar in (c) is 500 µm.

Rabies-GFP labeled neurons in the periaqueductal gray were predominately concentrated in the ventrolateral column. Case 6271 had many rabies-GFP labeled neurons in the medial paralemniscal nucleus, and all these neurons were mutually exclusive with the TVA-mCherry-expressing population of NPS neurons immediately dorsal to them in the semilunar nucleus (**Figure 11**). The midbrain reticular nucleus contained scattered rabies-GFP labeled neurons. Most of these neurons, like rabies-GFP labeled neurons in the pontine and medullary reticular formation, had prominent dendrites.

**Figure 11.**
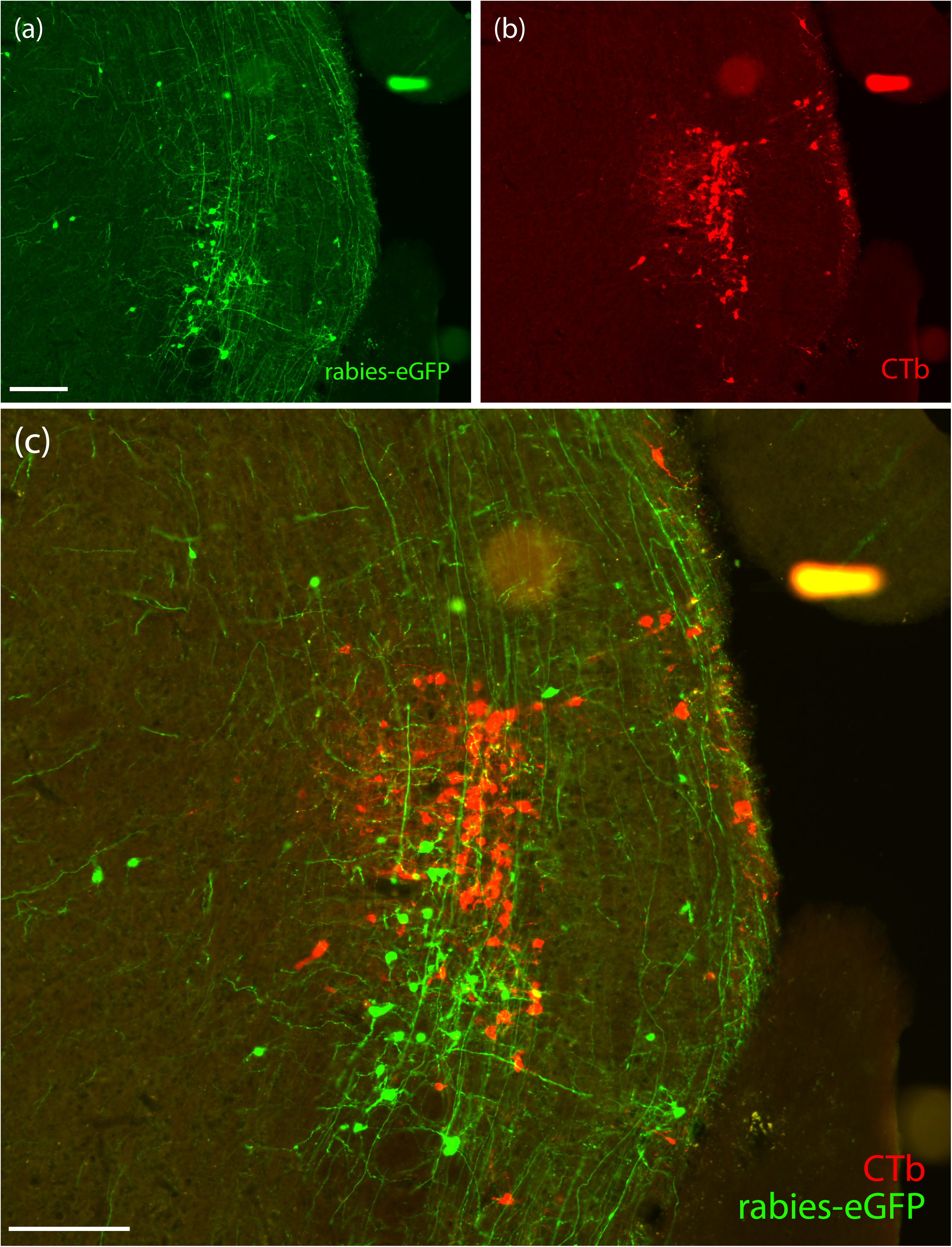
Rabies-eGFP-labeled neurons (green in a, c) in the medial parameniscal nucleus distributed ventral to TVA-mCherry labeled neurons (red in b, c) in the semilunar nucleus of case 6271. Scalebar in (a) is 200 µm and applies to (b). Scalebar in (c) is 200 µm.

### Hindbrain

Several auditory-related regions of the hindbrain were prominently labeled with rabies-GFP. The dorsal and ventral cochlear nuclei contained many rabies-GFP labeled neurons. In unilateral cases, the contralateral cochlear nucleus contained the vast majority of rabies-GFP labeled neurons (**Figure 10**). The superior olivary complex contained a moderate number of rabies-GFP labeled neurons, and in unilateral cases, most of these neurons were ipsilateral to the rabies starter cells (**Figure 10**).

The pontine reticular formation contained a moderate amount of dispersed rabies-GFP labeled neurons in every case. Few rabies-GFP labeled neurons were also found in the central pontine grey. The principal sensory trigeminal nucleus contained a few neurons in every case, and the spinal trigeminal nucleus contained a moderate number of rabies-GFP labeled neurons. We also found a significant number of rabies-GFP labeled neurons scattered throughout the medullary reticular formation (**Figure 10**). Small numbers of rabies-GFP labeled neurons were found in the nucleus of the solitary tract and area postrema, gracile nucleus, cuneate nucleus, and vestibular nucleus.

### Cerebellum

While the cerebellar cortex did not have any rabies-labeled neurons in most cases, we found one Purkinje neuron expressing GFP in the flocculus of case 6269. We found one or a few neurons were scattered in deep cerebellar nuclei of every case except 6272.

## Discussion

This paper builds upon our previous work on the molecular identity and efferent projections of NPS neurons. NPS neurons have a distinctive distribution, with a dense concentration in the extreme rostral-lateral part of the parabrachial region. From there, they extend rostrally into the semilunar nucleus and caudally into the dorsolateral parabrachial subnucleus. In contrast to the varied input patterns labeled by injections of the non-selective retrograde tracer CTb into this region, we found a consistent pattern of retrograde labeling in auditory-related regions and in the reticular formation after Cre-conditional rabies tracing. Below, we discuss these findings within the context of previous neuroanatomical studies, explore functional implications, and examine limitations of our approach.

### Comparison to previous neuroanatomical work

Input connections to NPS neurons in the parabrachial region have been examined by two groups investigating regulation of wakefulness (Angelakos et al., 2023; Xing et al., 2024). Similar to our approach, Angelakos et al. performed rabies retrograde tracing from the lateral parabrachial nucleus in *Nps*-IRES-Cre mice. They reported inputs from the bed nucleus of the stria terminalis, central amygdala, inferior colliculus, ventral cochlear nucleus, pontine reticular nucleus, and periaqueductal gray. However, they also reported that the second largest input in the brain arose from local neurons in the lateral parabrachial area (**Figure 2I** of Angelakos et al., 2023). In contrast, we found very few non-starter cells (rabies-eGFP neurons without TVA-mCherry) in the parabrachial region, likely because we allowed more time for TVA-mCherry expression, which enabled a clear distinction between starter cells (TVA-mCherry- and eGFP-expressing) and retrogradely labeled neurons (eGFP-only). Based on these findings, we conclude that local interneurons are not a major source of input to parabrachial NPS neurons. Besides this one exception, the proportions of inputs reported in that study are largely consistent with the pattern we found. Xing et al. also performed rabies retrograde tracing from the parabrachial region in *Nps*-IRES-Cre mice and reported inputs from the bed nucleus of the stria terminalis, central nucleus of the amygdala, and lateral hypothalamus. No injection site was shown, so the distribution of starter neurons was unclear, and the lack of labeling in auditory-related brain regions could be due to allowing only one week for TVA-mCherry expression before injecting rabies or due to the different helper viruses and modified rabies used in that study.

Our CTb retrograde tracing results were largely consistent with previous studies examining inputs to the parabrachial nucleus. Retrograde tracer injections in the parabrachial nucleus in rats labeled neurons in the medial prefrontal cortex and cortex overlying the claustrum, bed nucleus of the stria terminalis, substantia innominata, central nucleus of the amygdala, several hypothalamic nuclei, periaqueductal gray matter, ventral midbrain (Moga et al., 1990), as well as the hindbrain reticular formation, nucleus of the solitary tract, area postrema, and spinal trigeminal nucleus (Herbert et al., 1990). This pattern of retrograde labeling closely matches the pattern in our CTb case 7209, which had the most caudal injection site, spreading back into the middle of the parabrachial nucleus. Our CTb case 7216 lacked most of this pattern but had more prominent labeling in the ventrolateral medulla, likely due to the spread of this more rostral injection site ventrally, into the Kölliker-Fuse nucleus.

We are not aware of any previous studies with brain-wide analysis after retrograde tracer injections into the extreme rostral-lateral subregion that contains the dense concentration of NPS neurons, or of the rostral extension of this population into the semilunar nucleus. All of our CTb injections, with the exception of case 7209, had prominent labeling in auditory-related regions including the auditory cortex, inferior colliculus, nuclei of the lateral lemniscus, cochlear nucleus, and superior olivary complex. We were initially skeptical of these results because retrograde labeling in auditory brain regions might have been due CTb entering axons traveling through the lateral lemniscus (Chen & Aston-Jones, 1995), but the prominent rabies labeling in these same regions strongly supports an important role for auditory-related input to NPS neurons.

Input from auditory-related brain regions composed the plurality of all rabies-eGFP labeled neurons in every case. These regions corresponded to labeling in CTb cases with more rostral injection sites, with the notable exception of the auditory cortex. In contrast to the abundant retrograde labeling in the auditory cortex of every CTb case, we only found a few rabies-eGFP-labeled neurons in the cortex. This could reflect near-absence of direct cortical input to NPS neurons, but it may instead reflect a reluctance of rabies to label cortical neurons, as reported in previous rabies retrograde tracing studies (Sun et al., 2019; Ährlund-Richter et al., 2019). Channel rhodopsin-assisted circuit mapping (CRACM) from the auditory cortex in *Nps*-2A-Cre;R26-LSL-L10GFP Cre-reporter mice could help resolve this question (Petreanu et al., 2007).

### Functional implications

Neurons in the parabrachial nucleus relay sensory information including pain, temperature, taste, and visceral signals to the forebrain for behavioral responses (Carter et al., 2015; Chamberlin & Saper, 1992; Geerling et al., 2016; Jarvie et al., 2021; Kim et al., 2020; Nakamura & Morrison, 2008; Park et al., 2020). NPS neurons in the parabrachial region send projections to regions of the brain involved in stress and threat-response – paraventricular thalamus, dorsomedial hypothalamus, periaqueductal grey, and ventral bed nucleus of the stria terminalis – as well as regions implicated in circadian function: the subparaventricular zone, intergeniculate leaflet, and magnocellular subparafasicular nucleus (Zhang et al., 2024). Consistent with these projection patterns, previous studies explored a role for NPS neurons in increasing wakefulness (Angelakos et al., 2023; Xing et al., 2024). However, the function of NPS neurons in relation to their inputs remains mysterious.

The NPS neurons receive substantial auditory input, with lesser input from the reticular formation, BST, and CeA. The predominance of auditory-related input was particularly striking and unexpected. This strong input from neurons low in the auditory hierarchy suggests that NPS neurons receive minimally processed acoustic information. Beyond auditory inputs, the medullary and pontine reticular formation, followed by the spinal trigeminal nucleus, provided the next greatest inputs. The brainstem reticular formation contains diffuse isodendritic neurons that integrate heterogenous sensory information and help control posture, arousal, pain and startle responses, and cardiorespiratory control (FRENCH, 1958; Ramón-Moliner & Nauta, 1966; Schepens & Drew, 2004; Wang, 2009). The spinal trigeminal nucleus receives sensory information from the face, mouth, and other parts of the head directly from branches of the trigeminal nerve (Sessle, 2000). Additional input to NPS neurons comes from the BST and CeA, which are implicated in anxiogenic states and threat responses (Davis et al., 2010; Moscarello & Penzo, 2022; Walker et al., 2003).

Overall, the connectivity pattern of NPS neurons suggests that these neurons integrate anxiogenic and threat-related information from the BST and CeA with predominantly auditory information, supplemented by heterogeneous sensory information from the reticular formation and spinal trigeminal nucleus. This positions NPS neurons to assess and process auditory information in relation to threat and anxiety calculations and in the context of physiologically salient information. These neurons then relay this integrated information to forebrain areas that coordinate behavioral responses.

### Limitations

The Cre-conditional retrograde tracing approach we used begins with Cre-dependent expression of TVA-mCherry and G. Once activated by Cre, their expression and subsequent rabies-eGFP labeling is unrelated to the varying level of *Nps* expression in a given cell. This is a relevant consideration because NPS neurons in the main cluster within the extreme lateral parabrachial nucleus have consistently high levels of *Nps* mRNA and NPS immunoreactivity, while neurons in the semilunar nucleus and parabrachial dorsal lateral subnucleus express less *Nps* mRNA and often lack NPS immunoreactivity under baseline conditions (Huang et al., 2022). Rabies starter cells covered a wide distribution of *Nps*-expressing neurons in this region in our combined experiments, and retrograde labeling with rabies-eGFP presumably represents their combined total set of afferents. Exploring the possibility that distinct parabrachial and semilunar NPS subpopulations have distinct sets of afferent connections would require specific genetic markers to distinguish between them.

Additionally, rabies has decreased avidity for specific neuron types. After similar injection sites centered on the LHA, proportionally fewer neurons in the cerebral cortex were labeled by modified rabies than by retro-AAV (Sun et al., 2019). Rabies virus also may not label *Sst*-expressing interneurons in the prefrontal cortex, nonpeptidergic neurons in dorsal root ganglia, and parabrachial-projecting neurons in the caudal nucleus of the solitary tract (Albisetti et al., 2017; Korkutata et al., 2025; Ährlund-Richter et al., 2019).

Finally, because this technique produces equally robust eGFP expression in every labeled neuron, it is not useful for determining the strength of each input. Future anterograde tracing and CRACM experiments could address these limitations and determine the functional strength of individual connections.

## Conclusion

This study used conventional retrograde axonal tracing to identify the brain-wide pattern of afferents to the extreme rostral-lateral parabrachial region containing the densest concentration of NPS neurons. We then used rabies retrograde tracing to identify direct inputs to *Nps*-expressing neurons in this region. These findings reveal an unexpected role for NPS neurons as integrators of auditory information.

## Role of Authors

J.C.G. planned the project and secured funding. R.Z. and J.C.G. drafted and edited the manuscript. R.Z. performed stereotaxic injections, histology, and microscopy. R.Z. and J.C.G. drafted and edited the figures and figure legends.

## Acknowledgements

We thank Jon Resch for sharing the EnvA-G-deleted Rabies-eGFP and AAV8-CAG-FLEX-RabiesG viral vectors. Research reported in this publication was supported by the National Heart, Lung, and Blood Institute of the National Institutes of Health under Award Number T35HL166206. The content is solely the responsibility of the authors and does not necessarily represent the official views of the National Institutes of Health.

## Ethical approval

All procedures performed in studies involving animals were in accordance with the ethical standards of the institution or practice at which the studies were conducted.

## Data availability statement

The data that support the findings of this study are available from the corresponding author upon reasonable request.

